## SupplementaryFigures for "The Telomere Length Landscape of Prostate Cancer"

### Supplementary Figures

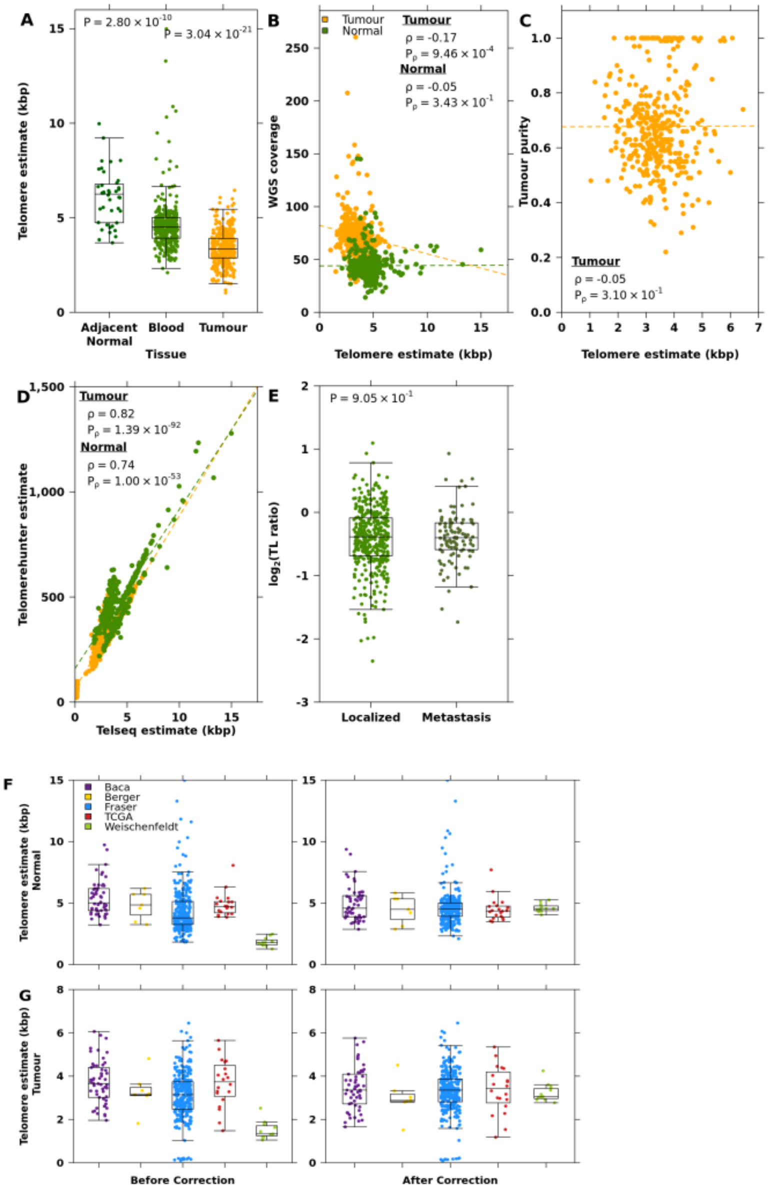


**Supplementary Figure 1 | Telomere length is independent of technical variables. A,** Comparison of TL in adjacent, histologically normal prostate tissue, blood and tumour tissue. *P* values are from a Mann Whitney U-test comparing adjacent normal TL to blood (left) and tumour TL (right). **B,** Correlation of telomere length (TL) estimated by TelSeq and WGS coverage **C,** tumour purity and **D,** TelomereHunter estimates. Green dots indicate non-tumour TLs, while orange dots indicate tumour TLs. Spearman's ρ and *P* values are displayed. **E,** Comparison of TL ratio in localized and metastatic prostate cancer samples. **F,** Non-tumour TL and **G,** tumour TL were batch corrected using a linear model (see **Methods**). Pre-corrected values are on the left and corrected values are on the right.

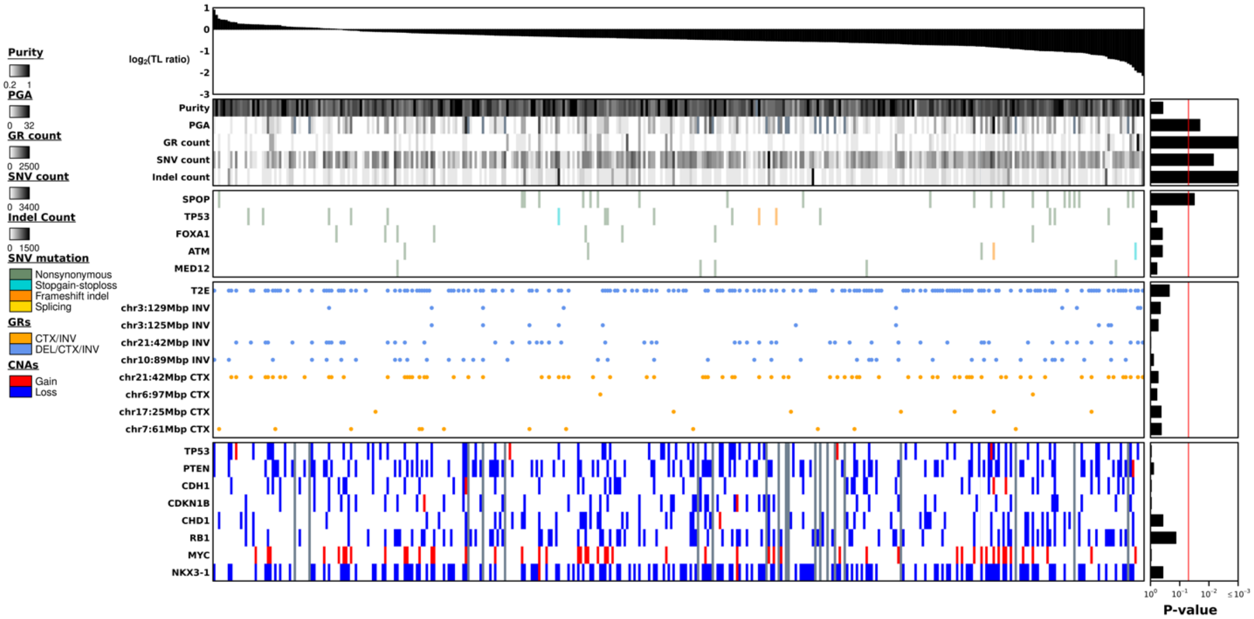


**Supplementary Figure 2 | Genomic associations with telomere length (TL) ratio**. TL ratio (tumour TL / non-tumour TL) is ranked in descending order. The association of TL ratio and measures of mutational burden, TMPRSS2:ERG (T2E) fusion status, as well as known prostate cancer genes with recurrent CNAs, coding SNVs, and GRs are shown. Bar plots to the right indicate the statistical significance of each association (see **Methods**).

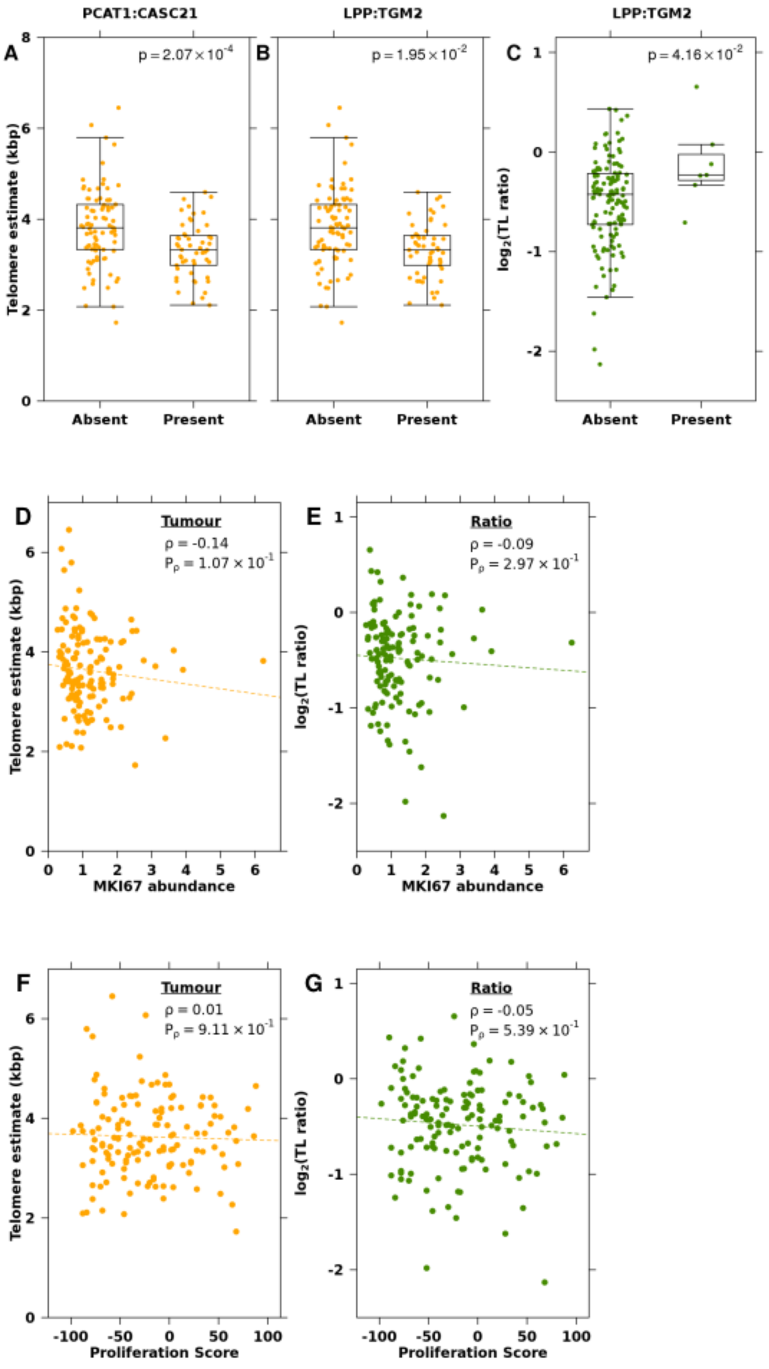


**Supplementary Figure 3 | Fusions are association with tumour TL and TL ratio. A,** Difference in tumour TL between samples with a PCAT1:CASC21 gene fusion and those without. **B-C,** Relationship of LPP:TGM2 fusions with **B,** tumour TL and **C,** TL ratio. *P* value from a Mann-Whitney U test is displayed. **D-E,** Correlation of MKI57 RNA abundance with **D,** tumour TL and **E,** TL ratio. **F-G,** Correlation of proliferation scores with **F,** tumour TL and **G,** TL ratio. Spearman's ρ and *P* values are displayed.

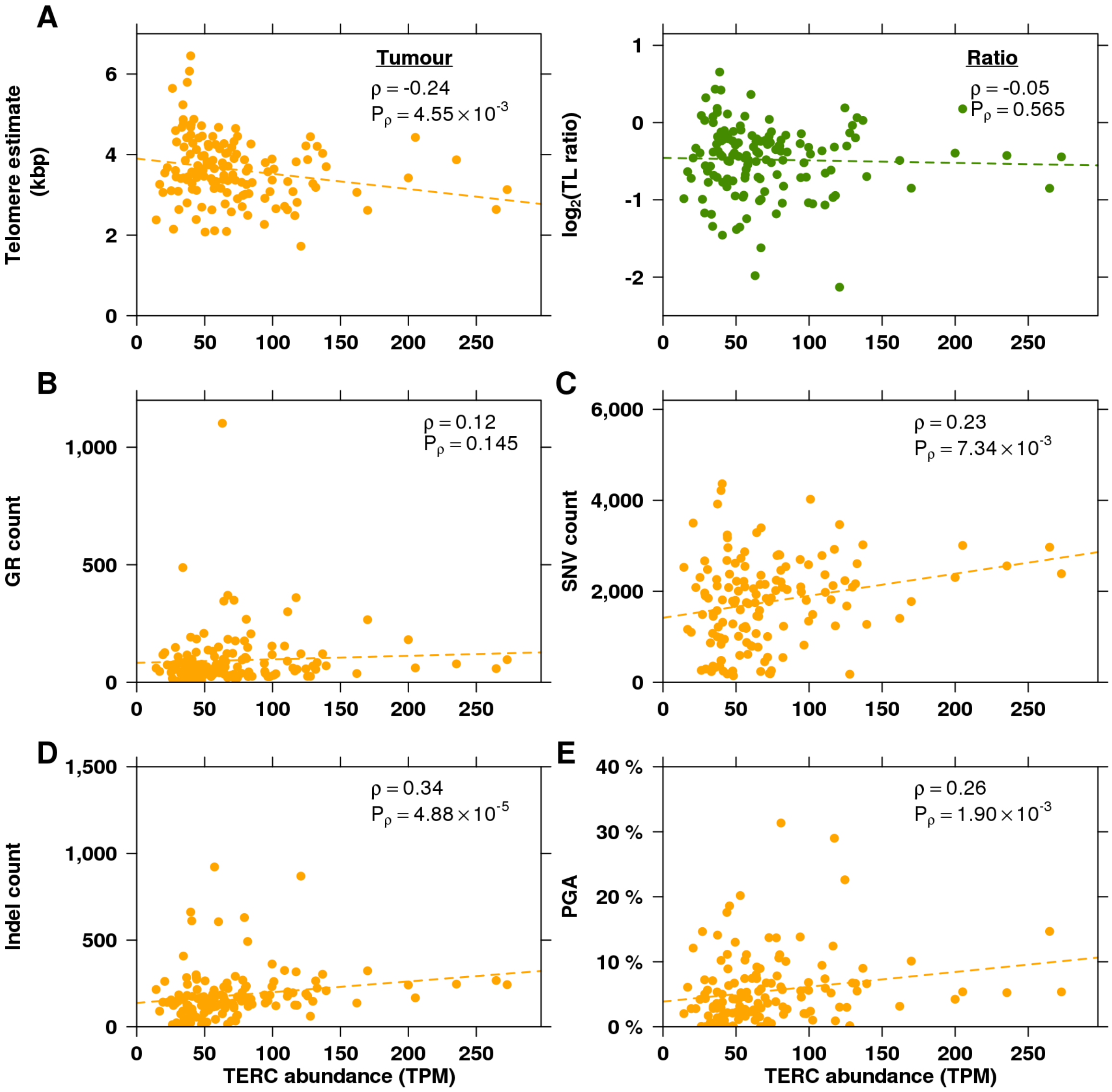


**Supplementary Figure 4 | Genomic correlates of TERC abundance. A,** Correlation of *TERC* abundance with tumour TL and TL ratio. Orange dots indicate tumour TL while green dots indicate TL ratio. **B-E,** Correlation of *TERC* abundance and the **B,** number of GRs, **C,** number of SNVs, **D,** number of indels and **E,** PGA. Spearman's ρ and *P* values are displayed.

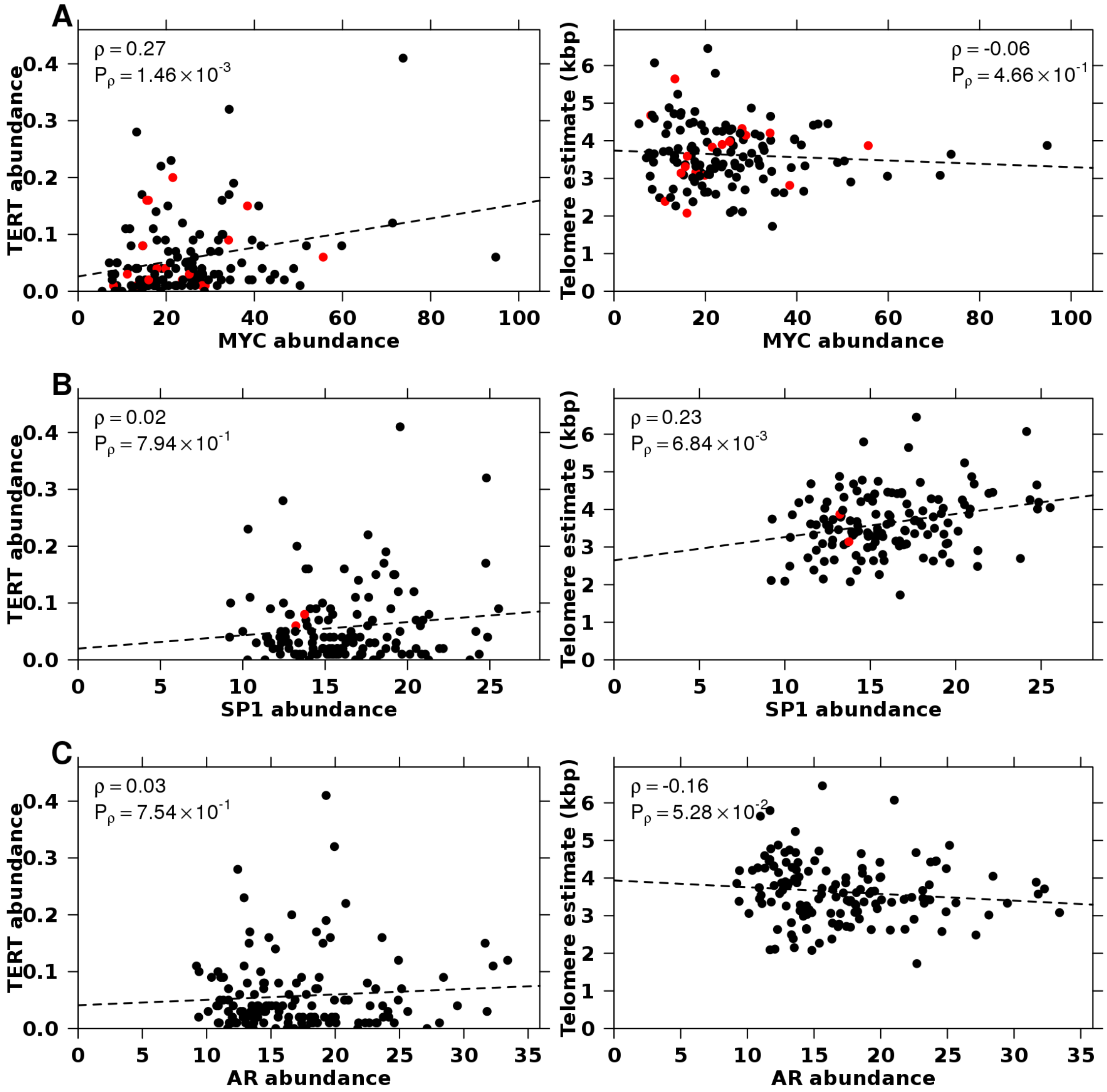


**Supplementary Figure 5 | Transcription factors of *TERT* are correlated with *TERT* abundance and telomere length (TL). A-C,** Association of *TERT* transcription factors **A,** *MYC,* **B,** *SP1* and **C,** *AR* with *TERT* abundance **(left panel)** and tumour TL **(right panel)**. Spearman's ρ and *P* values are displayed. Red dots indicate an amplification in that sample for the displayed transcription factor.

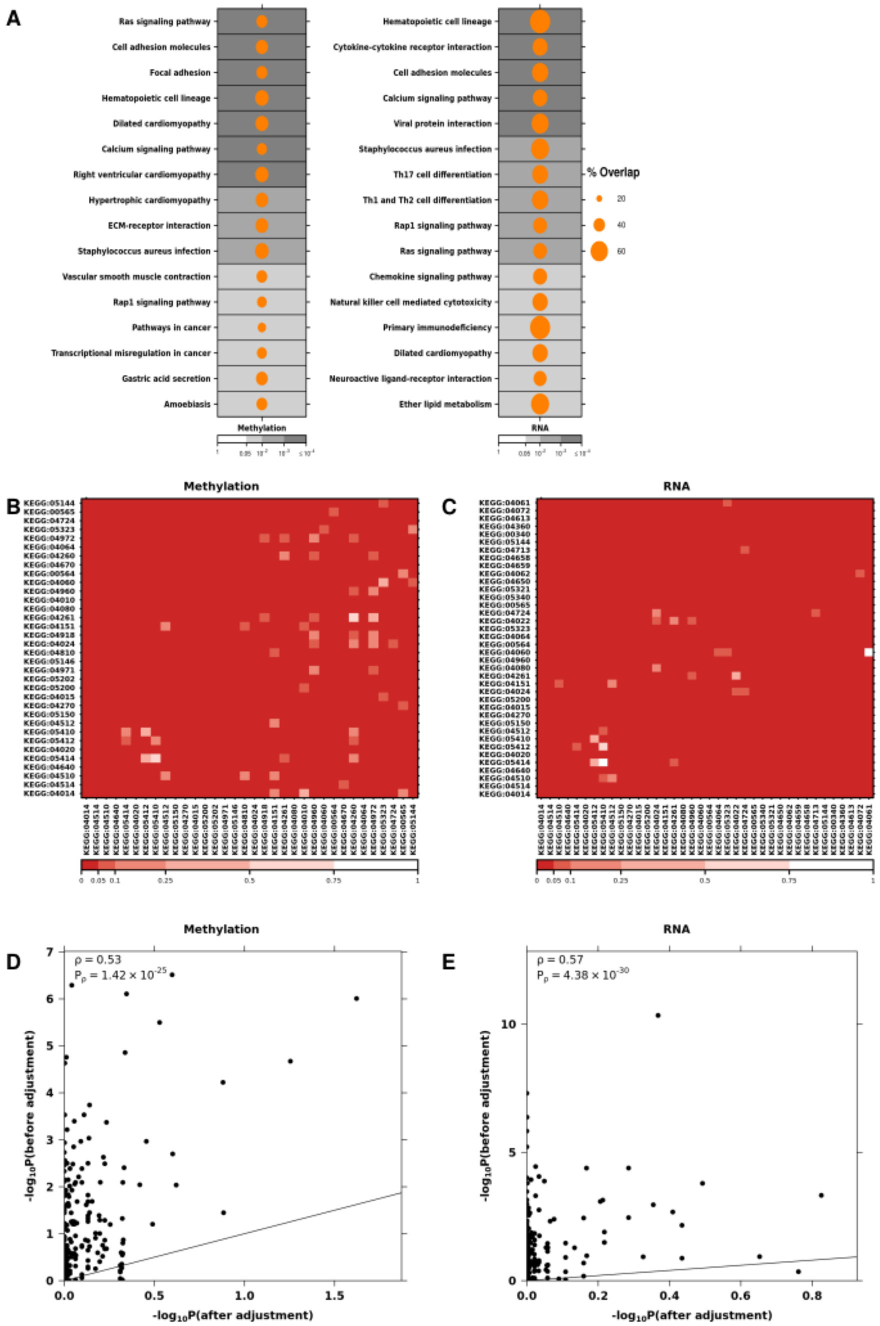


**Supplementary Figure 6 | Pathways enriched in genes with methylation or transcriptomic profiles that are correlated with tumour TL A,** Dotmap representing enriched pathways. Size of dot indicates the percentage of overlap between correlated genes and genes in the pathway. Background colour indicated unadjusted *P* values. **B-C**, Heatmaps of crosstalk matrices where white indicates loss of significance after removal of intersecting genes. **D-E,** Comparison of *P* values before and after crosstalk adjustment.

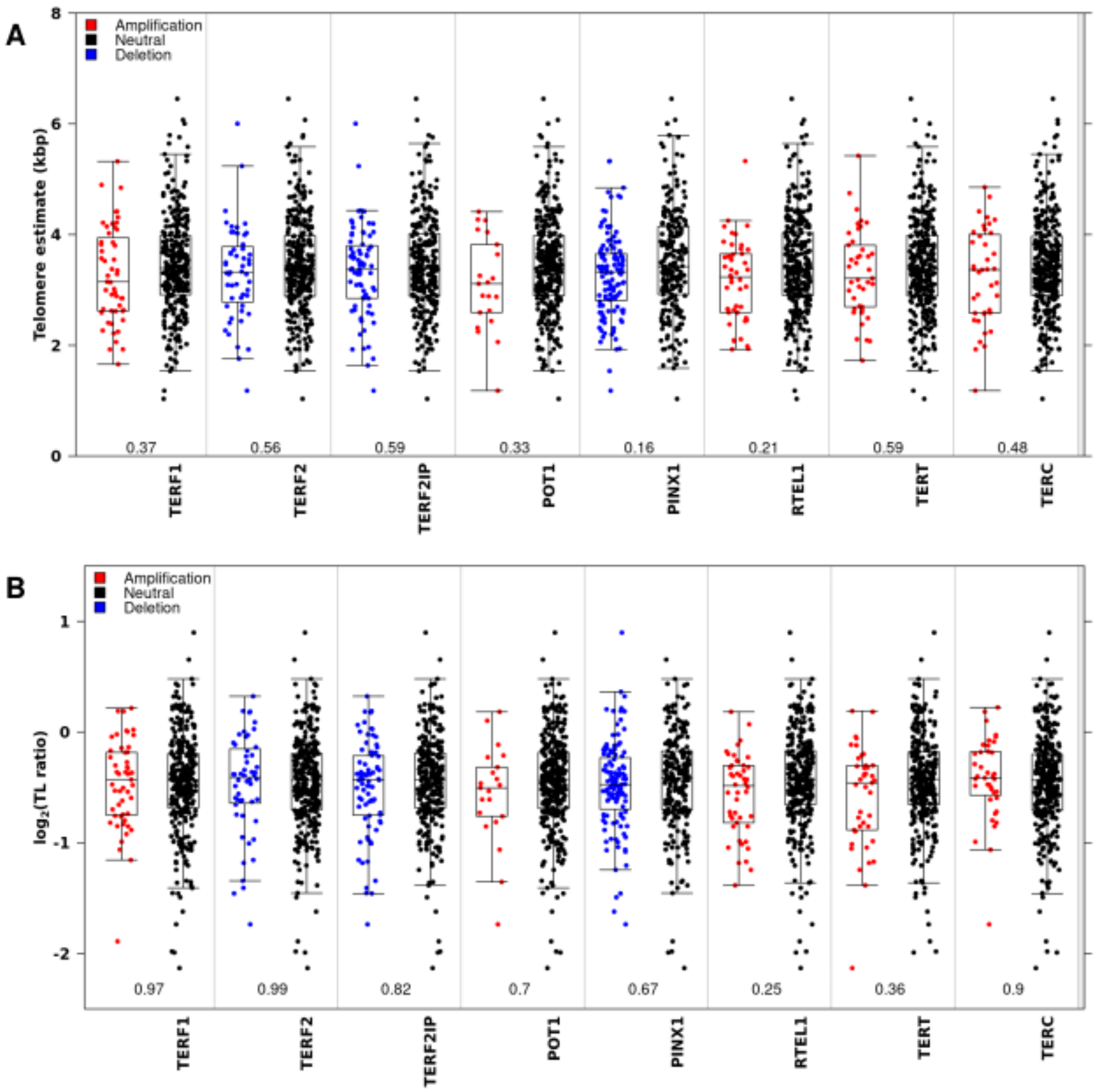


**Supplementary Figure 7 | Telomere length (TL) does not differ by copy number status in genes that make up the telomere complex. A,** Association between copy number aberrations in telomere complex genes and tumour TL. **B,** Association between copy number aberrations in telomere complex genes and TL ratio. FDR adjusted *P* values are from a Mann-Whitney U test. Colour of the dots indicate copy-number status of the gene: amplification (red), deletion (blue), or neutral (black).

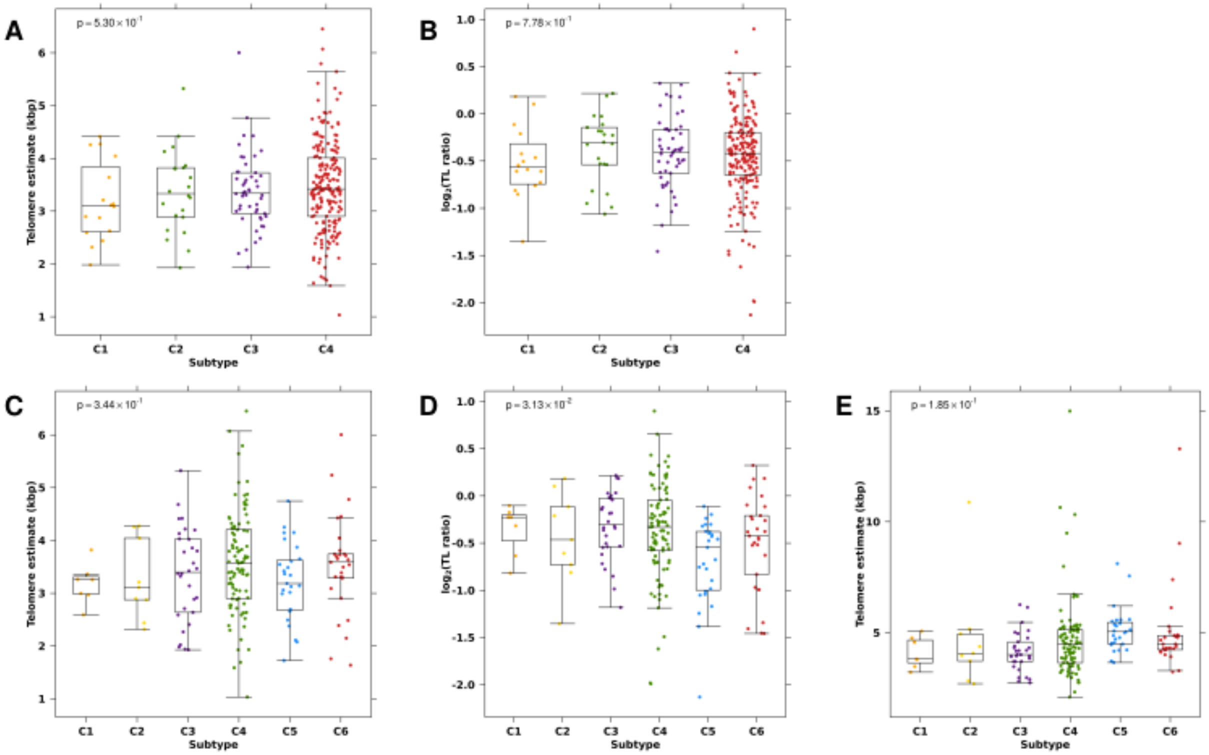


**Supplementary Figure 8 | Association of telomere length (TL) with CNA subtypes. A-B,** Association of **A,** tumour TL and **B,** $log_2$(TL ratio) with four previously identified CNA subtypes (Lalonde *et al*., 2014). **C-E,** Association of **C,** tumour TL and **D,** $log_2$(TL ratio) and **E,** non-tumour TL with seven previously identified CNA subtypes (Fraser *et al*., 2017).

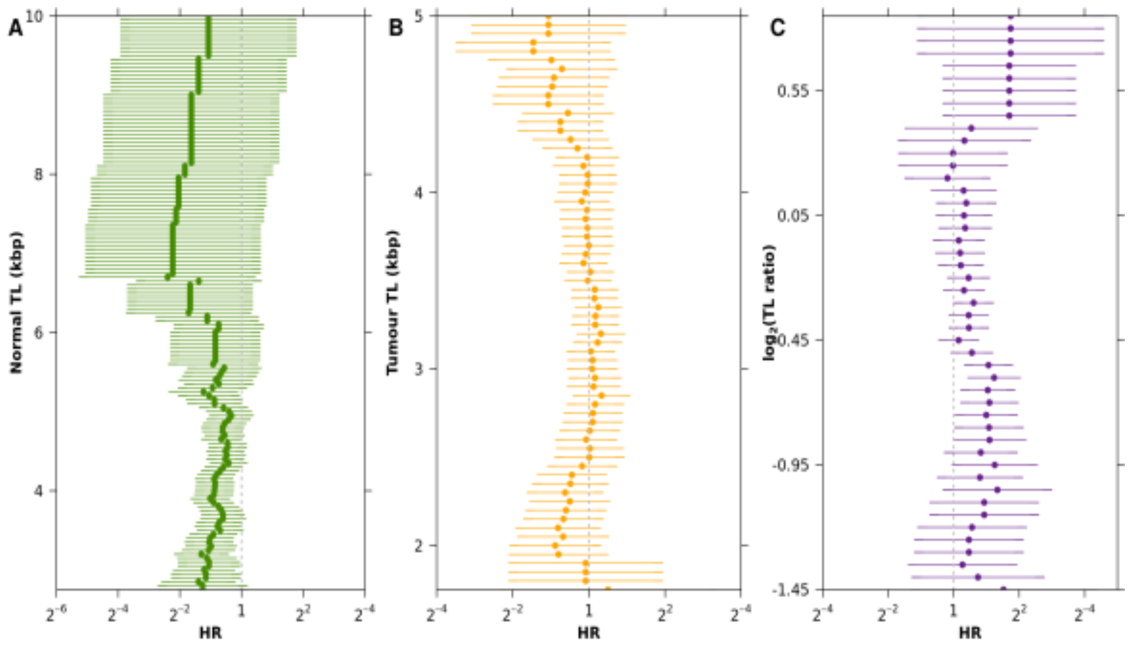


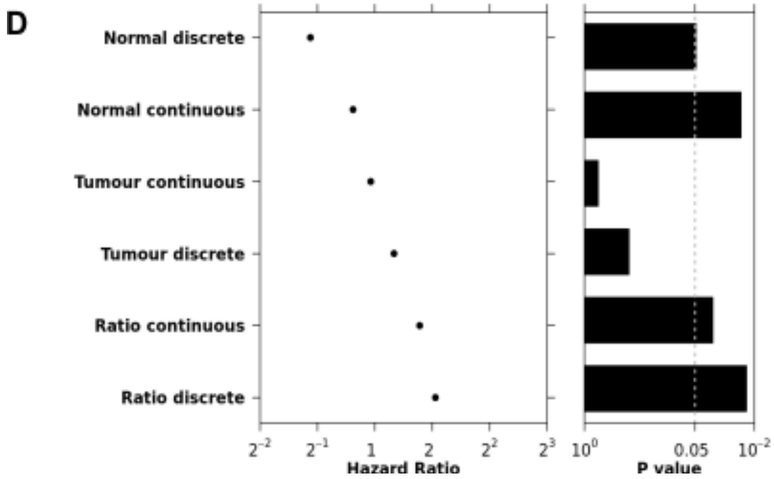


**Supplementary Figure 9 | Telomere length (TL) is associated with biochemical relapse (BCR). A-C,** Association of **A,** non-tumour TL, **B,** tumour TL and **C,** $\log_2$(TL ratio) with BCR using a Cox proportional hazards model at different TL cutoffs, incremented by 50 bp. **D,** Comparison of the best dichotomized Cox proportional hazards models and models fit with TL as a continuous value.
